# Bird populations most exposed to climate change are less responsive to climatic variation

**DOI:** 10.1101/2020.08.16.252379

**Authors:** Liam D. Bailey, Martijn van de Pol, Frank Adriaensen, Emilio Barba, Paul E. Bellamy, Jean-Charles Bouvier, Malcolm D. Burgess, Anne Charmantier, Camillo Cusimano, Blandine Doligez, Szymon M. Drobniak, Anna Dubiec, Marcel Eens, Tapio Eeva, Peter N. Ferns, Anne E. Goodenough, Ian R. Hartley, Shelley A. Hinsley, Elena Ivankina, Rimvydas Juskaitis, Bart Kempenaers, Anvar B. Kerimov, Anne Lauriere, Claire Lavigne, Agu Leivits, Mark C. Mainwaring, Erik Matthysen, Jan-Åke Nilsson, Markku Orell, Seppo Rytkönen, Juan Carlos Senar, Ben C. Sheldon, Alberto Sorace, Martyn J. Stenning, János Török, Emma Vatka, Stefan J.G. Vriend, Marcel E. Visser

## Abstract

The phenology of many species shows strong sensitivity to climate change; however, with few large scale intra-specific studies it is unclear how such sensitivity varies over a species’ range. We document large intra-specific variation in phenological sensitivity to temperature using laying date information from 67 populations of two European songbirds covering a large part of their breeding range. Populations inhabiting deciduous habitats showed stronger phenological sensitivity compared with those in evergreen and mixed habitats. Strikingly, however, the lowest sensitivity was seen in populations that had experienced the greatest change in climate. Therefore, we predict that the strongest phenological advancement will not occur in those populations with the highest sensitivity. Our results show that to effectively assess the impact of climate change on phenology across a species range it will be necessary to account for intra-specific variation in phenological sensitivity, climate change exposure, and the ecological characteristics of a population.

## Introduction

Environmental temperature is often an effective predictor of future conditions that impact organismal fitness ^1^. Because of this, many organisms exhibit a strong relationship between temperature and phenology leading to clear phenological advancement with anthropogenic climate change ^2–5^. The rate of phenological advancement is the product of a species’ ‘phenological sensitivity’ ^4^ and ‘climate change exposure’ (Box 1), which are both affected by the biotic and abiotic environment ^6–12^. For example, differences in the timing and availability of resources, due to factors such as habitat type, can affect both absolute phenology and phenological sensitivity ^6,11,12^. Similarly, climate change exposure can vary geographically, such as through the process of Arctic amplification where the rate of climate change is greater at higher latitudes ^8^. While it is well recognised that phenological sensitivity and climate change exposure can vary at an *inter*-specific level ^4,5,13^, it is poorly understood how *intra*-specific variance in these traits, and how they are correlated, may affect phenological advancement. Such analysis is important to determine the extent to which results from a single population are representative of a species across its range.

To quantify intra-specific variation in phenological sensitivity and climate change exposure we first need to identify the period during which temperature most strongly affects population phenology, termed a population’s ‘temperature window’ (Box 1). A population’s temperature window will directly determine the temperature values used to quantify phenological sensitivity and climate change exposure, yet in many populations we have little *a priori* knowledge on the temperature window used by the organisms under study ^4,14,15^. Quantifying phenological sensitivity using incorrect temperature windows can lead to an underestimation of sensitivity ^14^. Moreover, if appropriate temperature windows are used for some populations but not others, we may detect spurious intra-specific differences in phenological sensitivity. Similarly, using inappropriate temperature windows to calculate climate change exposure can lead to unreliable results due to the temporal heterogeneity of climate change intensity ^9,10^. Therefore, any attempt to understand intra-specific patterns of phenology must first account for potential intra-specific differences in temperature windows ^16–18^.

We use laying date information from 67 populations of two model bird species at a continental scale to quantify intra-specific variation in temperature windows, phenological sensitivity, climate change exposure, and their combined effects on phenological advancement (Box 1). We first quantify variation in the temperature windows of the two species across Europe and use these temperature windows to quantify population specific phenological sensitivity and climate change exposure. Next, we test potential biotic (habitat) and abiotic (photoperiod, precipitation) variables that may explain intra-specific variation in phenological sensitivity. Finally, we estimate how intra-specific variation in phenological sensitivity and climate change exposure affect a population’s phenological advancement. By focussing on two widely studied species we have the ability to obtain robust estimates of the drivers of intra-specific phenological variation that may help predict the effects of climate change on rare or under-studied species for which long-term multi-population data are unavailable.

#### Box 1: Key terminology and definitions used in this paper.

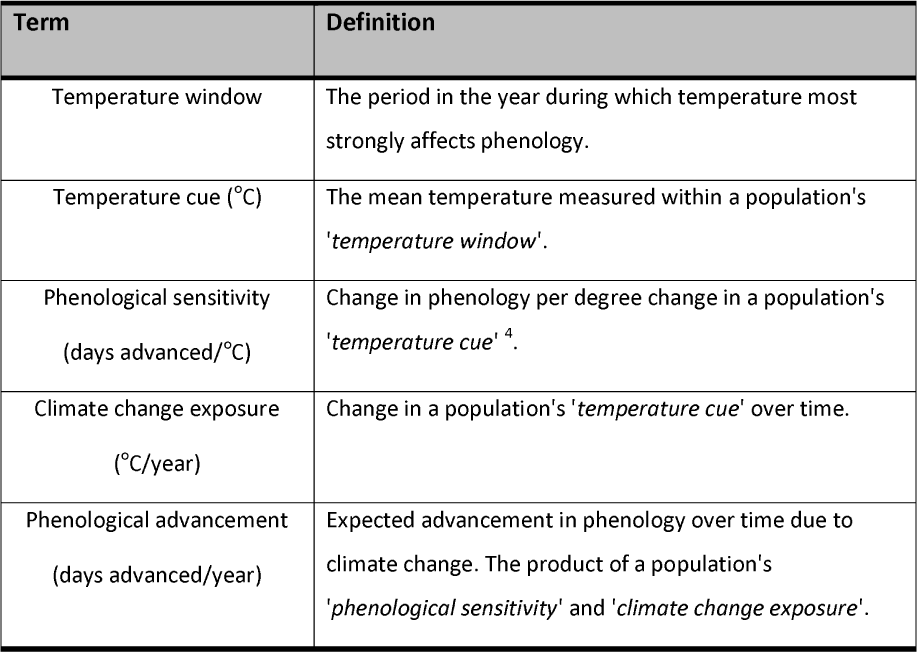

## Method

### Study populations

We collated data from 67 nest-box breeding populations of two closely related insectivorous passerines, the great tit (*Parus major)* and blue tit (*Cyanistes caeruleus*), across Europe (34 great tit and 33 blue tit; in 27 cases data for both species were collected from the same study site Fig. 1). We limited our analyses to populations for which a minimum of 9 years of data were available as we have previously been able to quantify temperature windows in a dataset of the same length ^19^. Sampled populations ranged latitudinally from 37.6° N (Italy) to 69.8° N (Finland), with the northern most populations close to the northern range limit of both species ^16,20^. Populations ranged in longitude from −3.99° W (UK) to 36.85° E (Russia). Populations were sampled from a range of habitats dominated by either deciduous or evergreen tree species or a mix of both.

**Figure 1:**
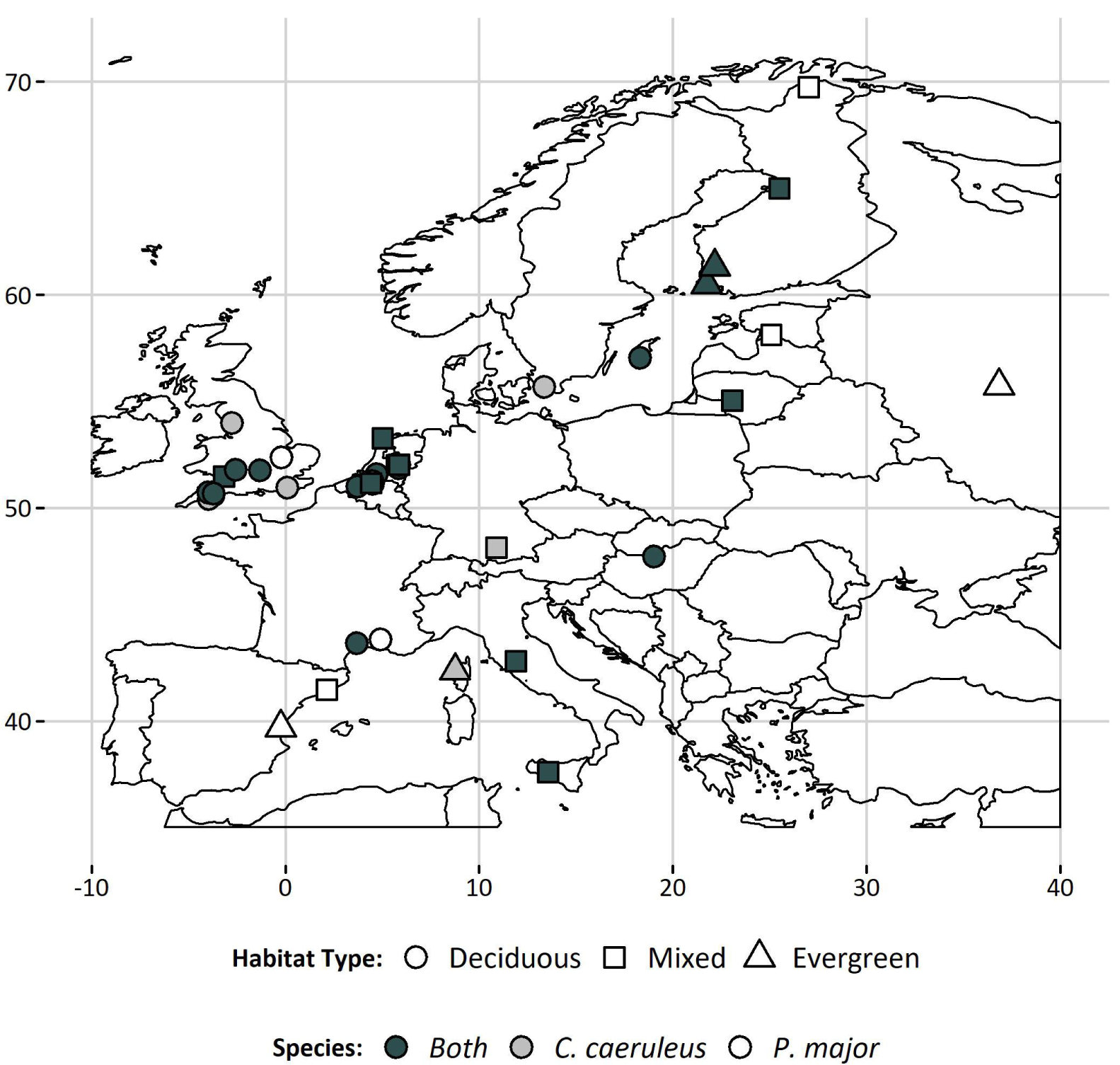
Distribution of study populations from which phenological data were collected. Phenological information (laying date) was recorded for both great tits (*Parus major*) and blue tits (*Cyanistes caeruleus*) at the majority of sites (68%; 27 sites).

### Phenological data

We quantified laying date (the date on which the first egg of a clutch was laid) for all females in all populations based on regular nest-box checks (at least weekly). When nests were not observed on the day the first egg was laid, laying date was estimated assuming one egg laid per day. For all analyses we used the laying date of first clutches and excluded second and replacement clutches. First clutches were defined as those laid within 30 days of the first clutch of the season in a given year and population ^21^. We used the mean annual laying date of first clutches as a measure of population phenology.

### Climate data

We extracted temperature data from the European Climate Assessment & Dataset E-OBS Gridded Dataset v17.0 ^22^. For every population, we extracted daily mean temperature (°C) for all years in which phenological data were available. In six populations, the study site location did not overlap with the gridded dataset. In four of these cases (Sagunto, Spain; Barcelona, Spain; Cardiff, UK; Askainen, Finland), we extracted temperature data from the nearest grid cell instead. The alternative grid cells were never more than 8km from the study site (3-8km). In one case (Vlieland, Netherlands) we interpolated daily mean temperature information from weather stations provided by the Royal Dutch Meteorological Institute (KNMI). All weather stations used to interpolate Vlieland data were from the neighbouring island of Terschelling (<33km away). As Vlieland is an island population, we considered local weather station data from a neighbouring island to be more reliable than the nearest grid cell located on the Dutch mainland. In the final case (Sicily, Italy), temperature data were taken from weather stations operated at the study site by a co-author (C. Cusimano).

### Estimating temperature windows

We determined temperature windows for each population using a systematic sliding window analysis in the R package ‘*climwin*’ ^*14,15*^. We followed the workflow described by van de Pol et al. ^15^ (their Fig. 3). As a first step, we designed the model to determine the relationship between daily mean temperature and mean laying date. For all populations, we used a linear model with a Gaussian error distribution with temperature as a fixed effect. Model residuals were weighted by the inverse of the standard error in annual mean laying date.

**Figure 2:**
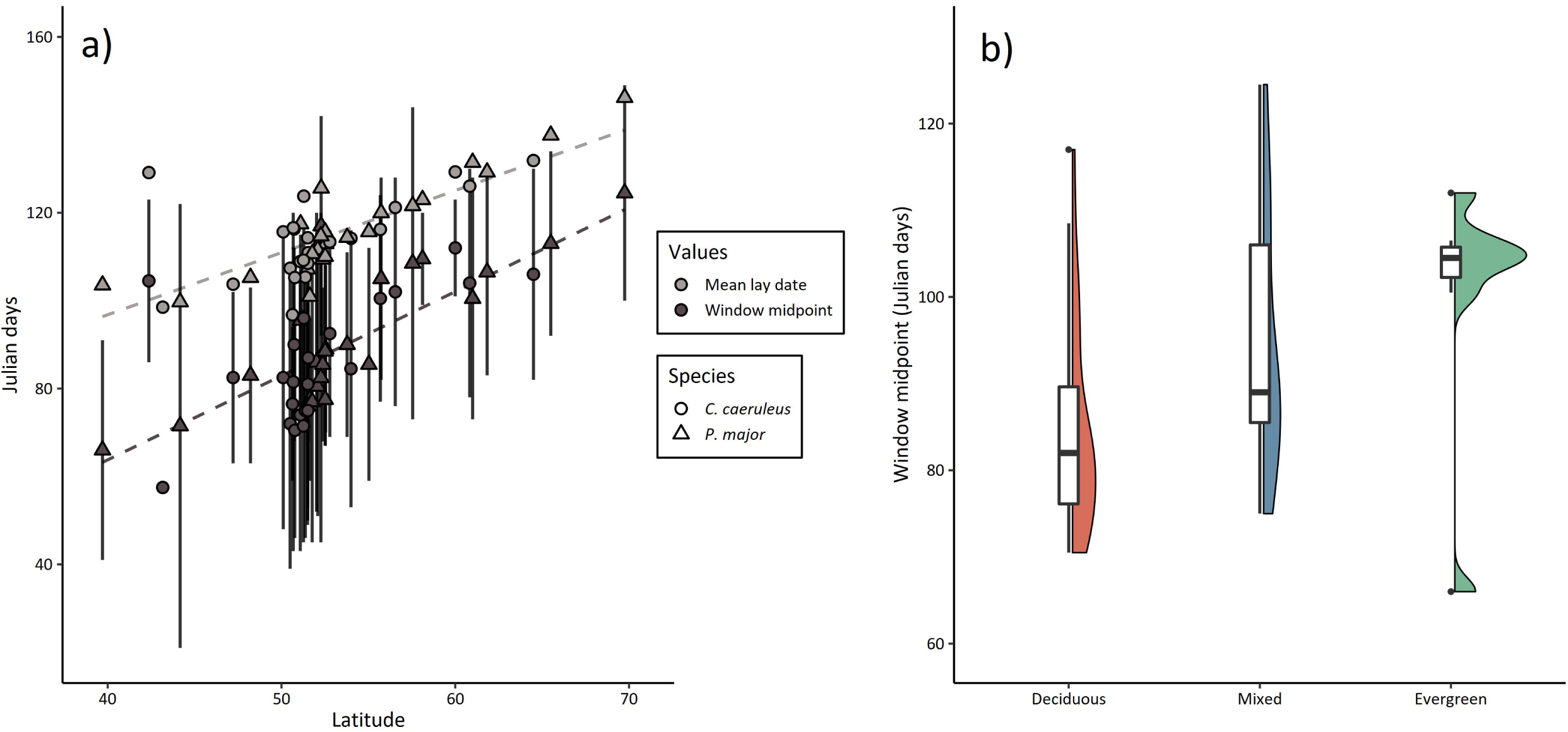
Intra-specific variation in temperature windows. a) Latitudinal change in temperature window midpoint (black) and mean annual laying date (grey) of European populations of great tits (triangles) and blue tits (circles). Temperature window midpoint increased with latitude in both species, while window duration (vertical bars) and the delay (difference between window midpoint and mean laying date; dashed lines) did not change significantly. b) Difference in temperature window midpoint for great and blue tits inhabiting deciduous (red), mixed (blue), or evergreen (green) habitats. Shown are box and violin plots of temperature window midpoints in each habitat type.

**Figure 3:**
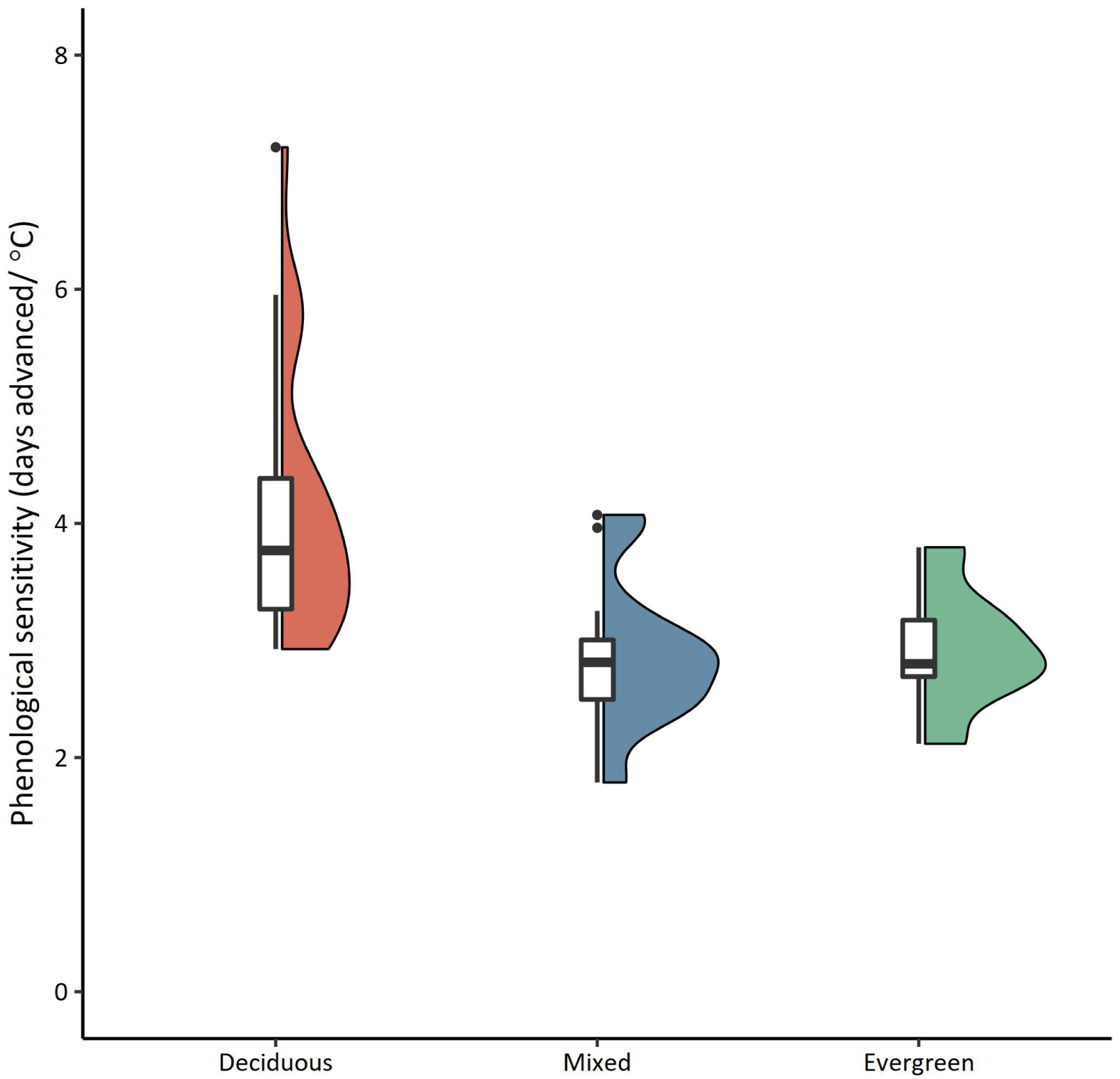
Variation in phenological sensitivity (days advanced/°C) between habitat types. Phenological sensitivity is higher in deciduous habitats than either evergreen or mixed habitats. Shown are box and violin plots of phenological sensitivity of great and blue tits in each habitat type.

We next selected relevant weather variables. Following previous studies on these species, our analysis focussed on mean temperature ^4,18,23^. We conducted all sliding window analyses using absolute windows (i.e. we assume that all individuals in a population had the same temperature window). We tested all potential temperature windows over a 365-day period before June 1^st^. For each potential temperature window, we estimated the relationship between mean temperature and mean annual laying date. Following previous studies, we tested for linear relationships between temperature and laying date ^4^.

As we tested a large number of potential temperature windows there were inherent risks associated with multiple-testing ^15^. To overcome this issue, we randomised the order of the original data in each population to remove any relationship between temperature and laying date and then re-ran the sliding window analysis. We replicated this randomisation procedure 100 times. We then compared our observed result to that of our 100 randomisations and determined the probability that our observed result could occur in a dataset where no relationship exists between temperature and laying date. This method is described in detail by van de Pol et al. ^15^. We used the metric P_ΔAICc_ for assessing the probability that the identified temperature window was a false positive ^14,15^.

We considered a temperature window to represent a true temperature cue if P_ΔAICc_ was ≤0.05 (i.e. the chance of such a result occurring in a randomised dataset was ≤5%). Populations with a best temperature window ≤14 days in duration were also excluded, as such short windows are biologically less plausible and can produce statistical artefacts ^15^. In all populations where we identified a true temperature cue (70%; see Results), we determined the duration and midpoint of the temperature window. Populations where false positive signals were identified were not used for further analyses. Temperature windows were detected more frequently in populations with more years of data (Fig. S1).

### Phenological sensitivity to temperature

Our sliding window method estimated the relationship between mean temperature and laying date in the best supported temperature window. However, a correlation between temperature and laying date may arise due to unmeasured, non-climatic variables that lead to a shared temporal trend between temperature and laying date ^24^. We used structural equation models to quantify the relationship between temperature and laying date after accounting for shared trends over time using the R package ‘*lavaan*’, using temperature data from population specific temperature windows. We used non-parametric bootstrapping with 1,000 iterations to estimate standard errors from these structural equation models.

### Intra-specific variation in temperature windows

To understand variation in the midpoint, duration, and delay of temperature windows (time lag between temperature window midpoint and mean laying date) we built general linear mixed effects models with a Gaussian error term. Population ID was included as a random intercept to account for cases where both species were sampled at the same site (Fig. 1). Population intercepts were assumed to be normally distributed with mean = 0 and variance = *σ*^2^. Our models included latitude, longitude, species, and habitat type (deciduous, evergreen, or mixed) as fixed effects, plus an interaction between species and latitude and longitude.

### Biotic and abiotic drivers of phenological sensitivity

After accounting for variation in temperature windows we next assessed which variables best explain phenological sensitivity in different populations. We fitted a general linear mixed effects model with a Gaussian error term with population ID included as a random intercept. We included a fixed effect to account for precipitation differences between populations, derived from the principal component analysis of Metzger et al. ^25^ which incorporates precipitation (mm) over the year (mean monthly precipitation January, April, July, October, and November). We had no clear expectation for the period during which precipitation should affect phenology, therefore we included this broad measure of precipitation patterns. We included an effect of habitat type (deciduous, mixed, or evergreen), and day length (minutes between sunrise and sunset on April 1^st^; https://www.esrl.noaa.gov/gmd/grad/solcalc/sunrise.html). Due to limited sample size, populations in broad-leafed (n = 2) and coniferous (n = 5) evergreen habitats were grouped together. We also included interactions between species and each of our fixed effects. Latitude and longitude were not included in these models as we were interested in identifying specific biotic and abiotic drivers of phenological sensitivity, rather than documenting spatial patterns.

Temperature differences between sites might also influence phenological sensitivity; however, average annual temperature and day length were strongly positively correlated (Pearson’s *r*: 0.88) making them impossible to disentangle in our analysis. Day length was retained because previous work indicates that differences in photosensitivity may be a driver of phenological differences among populations ^16,17^.

### Intra-specific variation in phenological advancement

We next calculated the expected phenological advancement of each population as the product of its phenological sensitivity and climate change exposure (Box 1). We estimated population specific climate change exposure since 1950 (the earliest year of data from the European Climate Assessment & Dataset E-OBS Gridded Dataset) within the population specific temperature windows identified above. Phenological advancement was only assessed in populations where a clear temperature window was identified and was not assessed for populations where no temperature data were available from the gridded dataset (Sicily, Italy and Vlieland, Netherlands).

### Confidence intervals

For all general linear mixed effects models, we calculated 95% confidence intervals using parametric bootstrapping with 2,000 iterations.

### Software

All analyses were conducted using R (v. 3.6.1) ^26^ in RStudio (v. 1.2.5019).

## Results

We identified temperature windows in 70% of our populations (24/34 great tit, 23/33 blue tit). Temperature windows were detected more frequently in populations with more years of data (Fig. S1), suggesting that failure to detect a temperature window after randomisation is due to limited sample size rather than phenological insensitivity. Laying date advanced with increased temperature in all 47 populations where we detected a temperature window (Data S1). None of the models showed evidence for significantly different responses between great and blue tits (Table S2).

### Intra-specific variation in temperature windows

There was a clear latitudinal pattern in temperature windows of both species, with populations at higher latitudes having temperature windows with midpoints later in the year (β = 1.70 days/degree latitude; 95% CI: 0.850/2.649; Fig. 2a; Table S3). Temperature window midpoint showed no clear relationship with longitude, nor was there any relationship between the temperature window duration and the latitude or longitude of populations in either species (Fig. S2; Table S3). The average delay between temperature window midpoint and mean laying date was 26.5 days (8.59 – 41.01) and showed significant no geographical pattern (Fig. S2; Table S3). Temperature windows were later in populations inhabiting evergreen forests compared to those in deciduous forests, although this difference could not be confidently distinguished from zero (β = 9.59 days; 95% CI: −2.50/21.98; Fig. 2b; Table S3).

### Drivers of phenological sensitivity

We observed a four-fold difference in the phenological sensitivity, ranging from 1.8 days/°C (Estonia) to 7.2 days/°C (Okehampton, UK; Fig. 3). Sensitivity was significantly associated with habitat type (deciduous, mixed, or evergreen) in both species, with significantly higher sensitivity in deciduous habitats than evergreen or mixed habitats. Phenological sensitivity was unaffected by day length, precipitation, or species (Table S4).

### Covariance between sensitivity and climate change exposure

Climate change exposure over the past seven decades varied from 0.01 °C/year (East Dartmoor, UK) to 0.05 °C/year (Upeglynis, Lithuania). Those populations with the highest phenological sensitivity tended to have experienced lower climate change exposure (Fig. 4; Data S1), with a negative correlation observed between phenological sensitivity and climate change exposure (Pearson’s *r*: −0.55). Expected phenological advancement, the product of sensitivity and exposure, showed a four-fold difference among populations (0.05 - 0.19 days advancement/year), a similar magnitude of variation to that seen in phenological sensitivity and climate change exposure. Great and blue tits in Gotland (Sweden) had the largest expected advancement, while those in East Dartmoor (UK) had the smallest.

**Figure 4:**
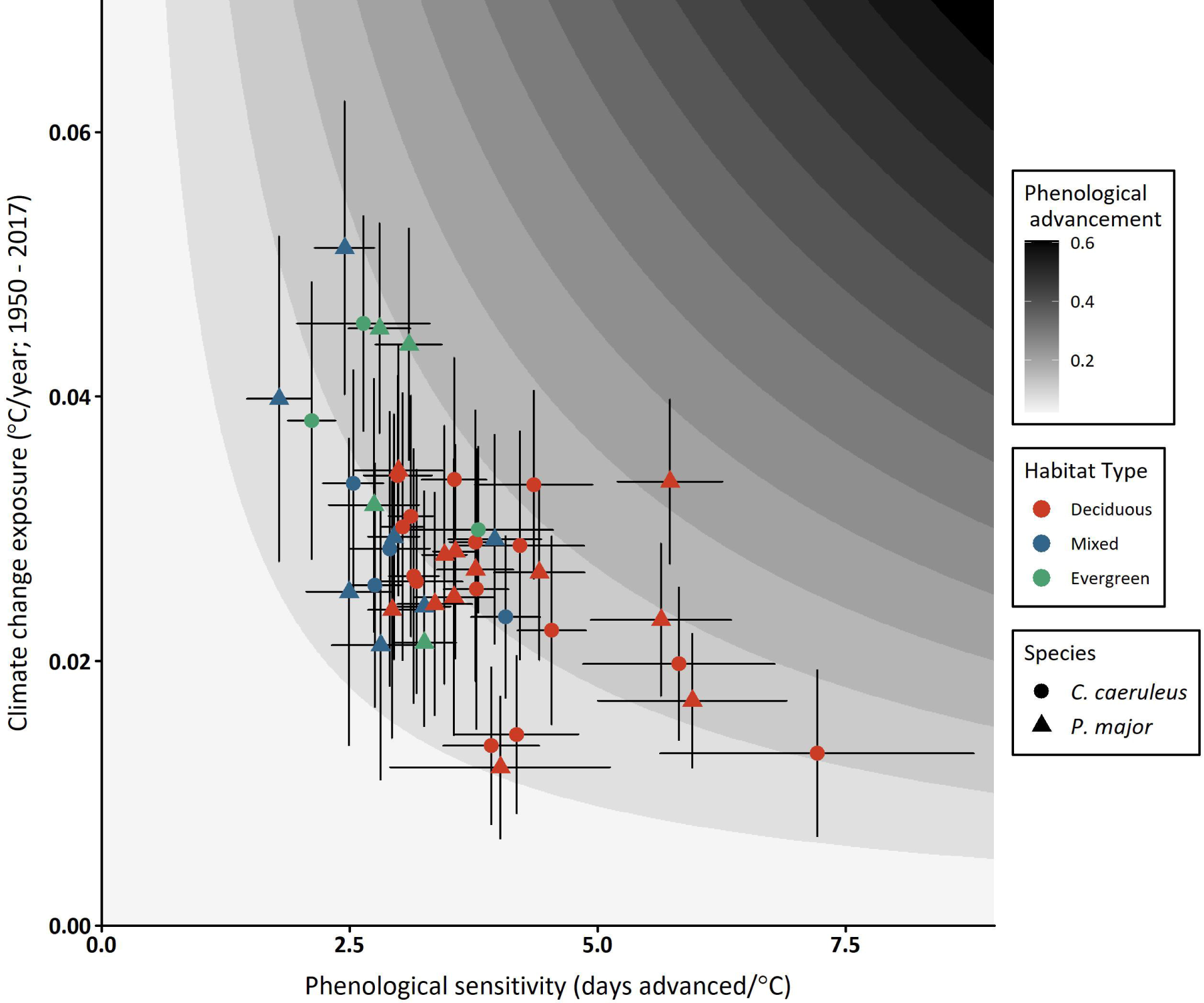
Estimated phenological advancement (days advanced/year) of great and blue tits across Europe. Phenological advancement (background colour) is a product of phenological sensitivity and climate change exposure. Populations with the highest recorded phenological sensitivity do not show the highest expected phenological advancement due to their lower climate change exposure. Note that phenological sensitivity and climate change exposure for each population are calculated within population specific temperature windows, which accounts for observed intra-specific variation in temperature window midpoints (Fig. 2).

## Discussion

We studied phenology in 67 populations of great and blue tits over a large part of their breeding range, within 47 of which we were able to identify temperature windows. In all 47 populations, earlier laying dates coincided with warmer conditions (Data S1). Phenology of populations varied spatially, with populations at higher latitudes showing later egg laying as well as later temperature windows. There was a four-fold difference in the strength of phenological sensitivity among populations. Differences in phenological sensitivity were associated with habitat type, with stronger sensitivity observed in deciduous dominated habitats. Interestingly, sensitivity was not an effective predictor of expected phenological advancement due to the covariance between sensitivity and climate change exposure. In both study species, those populations that showed stronger phenological sensitivity had experienced *less* warming over the past decades.

Our results demonstrate the potential for large intra-specific variation in both sensitivity to temperature and climate change exposure, which together can lead to intra-specific variation in the impacts of climate change. In fact, we detected no differences in phenological sensitivity between our two sympatric study species, which demonstrates that the ecological characteristics of a study population can be a better predictor of climate change effects than species traits, a pattern also observed in body condition responses to climate change ^27^. The presence of such intra-specific variation limits the efficacy of using single populations to draw conclusions about the impacts of climate change across a species’ range. Where possible, sampling from multiple populations with diverse ecological characteristics over a broad geographic scale should be undertaken to provide a better estimate of climate change impacts. In rare or poorly studies species, where such long-term multi-population data is unavailable, it will still be important to account for expected drivers of intra-specific variation when seeking to extrapolate from single populations. The results of this study provide an estimate of habitat type effects that will be relevant for such extrapolation, but it will be necessary to first understand how generalisable these results are in other species with different life history traits.

### Intra-specific variation in temperature windows

We used a standardised method to estimate temperature windows and show substantial intra-specific variation in temperature window characteristics. The most obvious pattern was that in latitude, with the phenology of birds at higher latitudes affected by temperature later in the year (Fig. 2). Change in the midpoint of temperature windows with latitude has now been clearly documented both here and in previous work ^18,23,28^, and is likely a consequence of differences in photoperiod thresholds and availability of food resources ^16^. The delay between a population’s temperature window and mean annual laying date did not vary spatially, suggesting that birds may require a fixed period of time to prepare for reproduction.

The factors that affect the duration of temperature windows are less clear. Previous work on great tits and pied flycatchers (*Ficedula hypoleuca*) in the UK ^18^ and Norway ^28^ documented shorter temperature windows at higher latitudes. However, both our study and multi-population work by Samplonius et al. ^23^ found no evidence of such pattern. We expect these studies would be able to detect any latitudinal effects, given that they covered a broad latitudinal range, so it seems likely that factors other than latitude may drive patterns in temperature window duration.

### Mechanisms driving phenological sensitivity

We observed large intra-specific differences in phenological sensitivity (Data S1); however, attributing a mechanism to the observed patterns must be done with caution. Phenological sensitivity at the population level may reflect individual level responses to temperature (i.e. phenotypic plasticity), but such patterns could also arise through other mechanisms. For example, if late nesting birds forgo reproduction in warmer years we would still detect a negative relationship between laying date and temperature without any phenotypic plasticity ^29^. Even if phenotypic plasticity explains the observed differences in sensitivity, we cannot tell whether these reflect intra-specific differences in the selective landscape or simply in the optimality of individual responses. It is possible that the optimal level of phenotypic sensitivity is similar across all study populations, but some populations have been able to track this optimum more closely ^30^.

As we carried out our analyses at the population level, we cannot definitively disentangle these mechanisms; however, previous work in a subset of our study populations has demonstrated that differences in phenological sensitivity between populations can be attributed to individual level differences ^12^ and that populations tend to show similarly optimal levels of phenological change even as the phenological optimum shifts across latitudes ^31^. With these previous results in mind, we expect that observed population level phenological sensitivity is likely driven by optimal shifts in individual phenology.

### Selective drivers of phenological sensitivity

The strength of phenological sensitivity was related to habitat type (Fig. 3), which confirms results from previous analyses conducted at a smaller spatial scale ^12^. Interestingly, this observed habitat effect holds over a broad latitudinal range, with low phenological sensitivity observed in both Fennoscandian coniferous evergreen habitats (e.g. Askainen, Finland) as well as Mediterranean broad-leafed evergreen habitats (e.g. Corsica, France) (Data S1). Our results show that habitat type can not only impact absolute phenological values ^7,34,35^ but can also affect phenological sensitivity. The fact that this pattern is observed in both coniferous and broad-leafed evergreen habitats suggests that this relationship is driven more by differences in the characteristics of the ecosystem rather than the exact species composition.

Differences in food resources between habitat types may help explain the observed sensitivity patterns. Temperature is a laying date cue in great and blue tits ^35^, enabling breeding birds to synchronise peak offspring provisioning with the peak in caterpillar abundance. Evergreen habitats tend to have lower overall caterpillar abundance than deciduous systems ^34,36^, which may necessitate greater dietary flexibility in nestlings and reduce the reliance of breeding birds on caterpillars and corresponding temperature cues ^37^. Dietary differences between habitat types have been documented in Mediterranean great and blue tits ^36,38^, and this pattern may apply more broadly across Europe. Evergreen forests also tend to have a wider period of peak caterpillar abundance than deciduous systems ^34^, which may affect the selection landscape. A narrow caterpillar peak in deciduous habitats will lead to a high fitness cost of mistiming ^39–42^, creating strong selective pressure for synchronisation by tracking temperature cues. In comparison, a broad caterpillar peak in evergreen habitats will reduce the costs of asynchrony, leaving birds less strongly temperature constrained. Finally, temperature cues have been shown to provide a less reliable indicator of caterpillar abundance in evergreen than deciduous habitats ^12^, which may lead populations to rely on alternative cues, such as vegetation phenology ^33^. All these factors may explain the lower phenological sensitivity in evergreen forests, but a European wide comparison of food resources available to breeding birds and their temporal dynamics would be necessary to understand this further.

If observed habitat patterns are a consequence of resource availability, our results may also be applicable outside our study species. We predict that other insectivorous passerines that use caterpillars as a primary food resource during offspring provisioning will show similar differences in phenological sensitivity between habitat types, although this will likely depend on the species’ dietary specialisation. However, these patterns might even extend beyond this specific feeding guild. For example, a relationship between patterns of resource availability and phenology has also been proposed for breeding shorebirds in Greenland ^11^, suggesting that the importance of food peak structure on phenology may be broadly generalisable to any species that relies on a seasonal peak in resources and is therefore vulnerable to phenological mismatch. Some examples may include invertebrates reliant on budding vegetation ^43^, secondary and tertiary consumers that utilise peaks in juvenile prey ^43^ or migratory arrivals ^44^, and plants reliant on insect pollinators ^45^. If habitats differ in the abundance, predictability, or the length of availability in these seasonal resources this may drive intra-specific variation in phenological sensitivity. In general, we predict that species that show dietary specialisation and live in habitats with narrow resource peaks will be more sensitive.

Although we show a clear effect of habitat type, it is important to note that we still observe variation in phenological sensitivity among deciduous and evergreen populations (Fig. 4). Some of this unexplained variation may be accounted for with more detailed estimations of habitat type that can differentiate deciduous and evergreen populations (e.g. proportion of deciduous trees, dominant tree species). It is also likely that there are other biotic or abiotic drivers of phenological sensitivity not accounted for here. This study provides a clear example of how the ecological characteristics of a population can affect sensitivity to climate change, and there is now a need for future work that can identify other potential drivers of intra-specific variation.

### Covariance between sensitivity and exposure

Identifying differences in sensitivity is only one component necessary to understand the phenological consequences of climate change, which will be the product of both a population’s sensitivity and climate change exposure (Box 1). The combined importance of sensitivity and exposure has been discussed extensively in an *inter*-specific context ^46–48^, but we show here that *intra*-specific (co)variance in these traits is also important. Spatio-temporal differences in climate change intensity can lead to differences in climate change exposure between populations from different geographic regions ^8^ or population’s with different temperature windows (Box 1) ^10^. While populations of great and blue tits inhabiting deciduous habitats showed strong sensitivity, many areas where deciduous populations are monitored (e.g. UK, Netherlands) have experienced limited climate change exposure. In contrast, higher latitude populations have experienced high climate change exposure over the past decades due to Arctic amplification ^8^, yet many of these populations reside in evergreen habitats and so have weaker sensitivity. In our study, the population with the highest predicted phenological advancement (Gotland, Sweden) was the most northerly deciduous population studied.

Quantifying the covariance between sensitivity and climate change exposure will be necessary to effectively predict which populations of a species will be most affected by future climate change. In our study, covariance between sensitivity and exposure made it unreliable to predict phenological advancement from either variable alone, both were needed to accurately predict the impact of climate change. Intuitively one might expect that a negative covariance between sensitivity and exposure would also buffer the intra-specific variation in phenological advancement. Although in many cases negative covariance is expected to reduce intra-specific variation, the reduction in our study was fairly small (∼24%; Supplementary material 2), and, as a consequence, intra-specific variation in phenological advancement was of a similar magnitude to variation observed in phenological sensitivity.

In general, it is still unclear whether covariance patterns between exposure and sensitivity are common and what sign they may have. Negative covariance between sensitivity and exposure may be particular to our study of phenology where both climate change exposure and the occurrence of evergreen habitats associated with lower sensitivity were affected by latitude. In other study systems, or when studying other traits, the sign of covariance may differ if sensitivity and exposure are affected by different drivers. Positive covariance could exacerbate existing patterns if the most sensitive populations are also those most exposed to climate change. Systems may also lack covariance between sensitivity and exposure in species that cover a limited geographic or ecological range.

Patterns of covariance could also shift over time due to concurrent changes in the biotic or abiotic environment. For example, climate change may alter plant communities through shifts in temperature and rainfall ^49^ or effects on plant pathogens ^50^. If climate change alters the ecosystem dynamics of a habitat (e.g. evergreen to deciduous dominant) we would expect knock-on effects on phenological sensitivity and phenological advancement. Therefore, our ability to predict impacts of climate change in populations will depend not only on our ability to quantify covariance between sensitivity and exposure, but also to predict future changes in those environmental characteristics that may affect sensitivity.

### Conclusions

To understand how populations and species will be affected by climate change, it is important to consider both their sensitivity to temperature change as well as their exposure to temperature shifts. This is often considered at an *inter*-specific level, but we show here that *intra*-specific variation in sensitivity and exposure also occurs and can influence how strongly populations will be affected by climate change. When seeking to understand climate change impacts, we should attempt to sample across a range of populations that experience different biotic and abiotic conditions to account for intra-specific patterns of (co)variance in sensitivity and climate change exposure. Without accounting for such intra*-*specific variation, we risk drawing conclusions from populations that are not representative of the species across its range.

## Acknowledgements

We would like to give a special acknowledgement to all the fieldworkers who have been invaluable in helping collect these many decades of data. We acknowledge the E-OBS dataset from the EU-FP6 project ENSEMBLES (http://ensembles-eu.metoffice.com) and the data providers in the ECA&D project (http://www.ecad.eu). Field study in Moscow region (EI & ABK) was supported by Russian Science Foundation (RSF Grant No. 20-44-01005). This study was also funded by research project CGL-2016-79568-C3-3-P (to JCS) from the Ministry of Economy and Competitivity, Spanish Research Council. AC was funded by the European Research Council (Starting grant ERC-2013-StG-337365-SHE). JT was funded by the Hungarian National Research, Development and Innovation Office (K-115970).

