## Supplementary material 1 for "Bird populations most exposed to climate change are less responsive to climatic variation"

**Supplement Material 1: Supplementary tables, figures, and data**

**Supplementary data S1:** Location, temperature window characteristics, phenological sensitivity, climate change exposure, and expected phenological advancement for all studied populations. Those populations where P_ΔAICc_ was ≤0.05 and top windows were > 14 days are shaded in grey (47 populations). For all other populations we could not confidently exclude the identified temperature window from the effects of over-fitting. These populations were excluded from further analysis. Estimates of phenological sensitivity, intercepts, and effect of year are determined using a structural equation model that accounts for potential shared trends between temperature and phenology. Note that the units of these coefficients are in negative April days, where days *before* March 31^st^ are positive and those *after* March 31^st^ are negative. This was done so that more sensitive populations have a more positive slope. Climate change exposure and expected phenological advancement are only included for those 47 populations where temperature windows could be confidently differentiated from effects of over-fitting. Expected phenological advancement is the product of phenological sensitivity and climate change exposure.

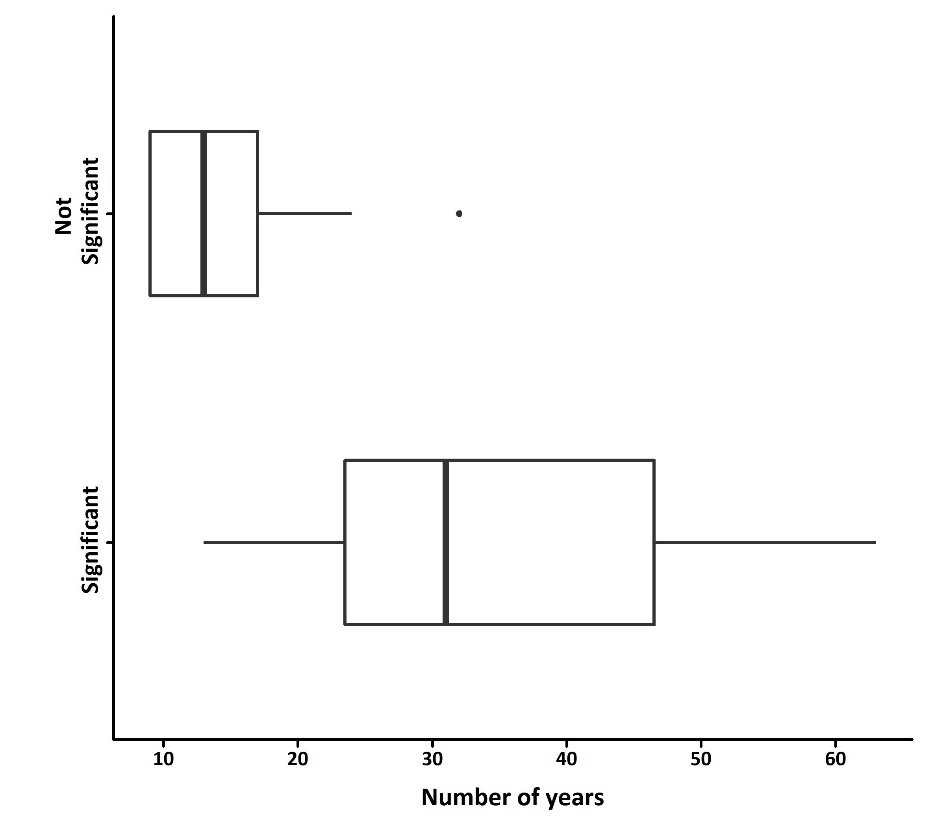

**Figure S1: Relationship between population sample size (number of years) and temperature window detection.** Temperature windows were identified using a cut-off of P_AICc_ ≤ 0.05. Those populations with temperature windows ≤ 14 days were excluded. Populations with no clear temperature window had fewer years of data than those where a temperature window was detected. Temperature windows will be more difficult to identify in datasets with limited sample size ^15^. Those cases without a clear temperature window are therefore likely a consequence of limited statistical power.

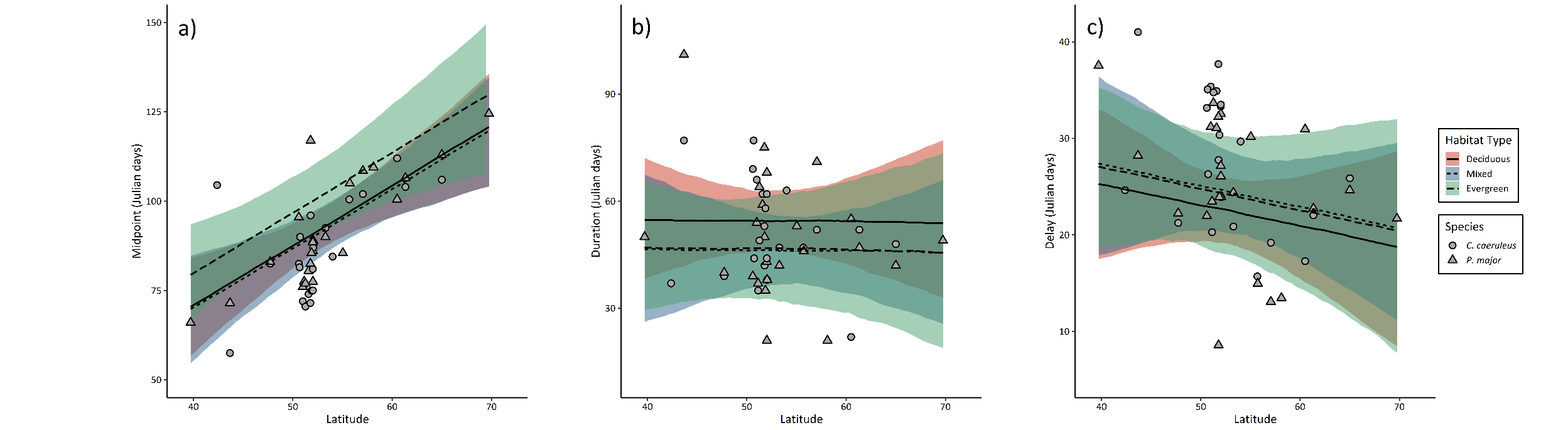

**Figure S2: Relationship between latitude and temperature window characteristics.** Change in temperature window a) midpoint, b) duration, and c) delay between midpoint and mean annual laying date in deciduous (red), mixed (blue), and evergreen (green) habitats. There is a clear relationship between latitude and temperature window midpoint. Temperature window duration and delay are unaffected. Plots show model predictions and 95% prediction intervals.

**Table S2:** Likelihood-ratio test results comparing models with and without species interactions. Coefficient estimates of all models with interactions are provided in Table S5 and Table S6.

| **Response variable** | **Log-likelihood (interaction model)** | **Log-likelihood (non-interaction model)** | **χ2** | **Degrees of freedom** | **p-value** |
| --- | --- | --- | --- | --- | --- |
| Midpoint | -163 | -166 | 5.88 | 2 | 0.053 |
| Duration | -190 | -191 | 1.99 | 2 | 0.370 |
| Delay | -151 | -148 | 5.10 | 2 | 0.078 |
| Phenological sensitivity | -44.3 | -45 | 1.42 | 2 | 0.840 |

**Table S3:** Effect of latitude, longitude, habitat type (deciduous, mixed, evergreen), and species (great or blue tit) on the midpoint (Julian day), duration (days), and delay (days) of temperature windows during which temperature most strongly affects laying date. Significant terms (where 95% confidence intervals do not overlap with 0) are shown in bold. The blue tit is used as the reference category for the species term. Deciduous is used as the reference category for the habitat type term.

| **Predictor variable** | **Parameter estimate  [95% confidence interval]** |
| --- | --- |
| **Midpoint** |  |
| Intercept | -3.319 [-49.574 / 39.04] |
| **Latitude** | **1.699 [0.850 / 2.649]** |
| Longitude | 0.091 [-0.454 / 0.604] |
| **Species (Great tit)** | **4.936 [1.722 / 8.279]** |
| Habitat (Evergreen) | 9.585 [-2.501 / 21.977] |
| Habitat (Mixed) | -0.718 [-10.096 / 8.095] |
| **Duration** |  |
| Intercept | 56.616 [-7.163 / 114.192] |
| Latitude | -0.001 [-1.129 / 1.253] |
| Longitude | -0.264 [-1.005 / 0.451] |
| Species (Great tit) | 0.638 [-6.886 / 8.464] |
| Habitat (Evergreen) | -7.863 [-24.157 / 8.902] |
| Habitat (Mixed) | -8.636 [-21.672 / 3.275] |
| **Delay** |  |
| **Intercept** | **38.636 [10.765 / 64.021]** |
| Latitude | -0.213 [-0.727 / 0.342] |
| Longitude | -0.281 [-0.609 / 0.035] |
| Species (Great tit) | -2.346 [-5.407 / 0.876] |
| Habitat (Evergreen) | 1.757 [-5.416 / 9.223] |
| Habitat (Mixed) | 1.82 [-3.959 / 7.047] |

**Table S4:** Effect of species (great or blue tit), precipitation, habitat type (deciduous, mixed, evergreen) and day length (minutes) on the strength of phenological sensitivity (days/^o^C). Significant terms (where 95% confidence intervals do not overlap with 0) are shown in bold. The blue tit is used as the reference category for the species term and deciduous habitat is the reference category for habitat type.

| **Predictor variable** | **Parameter estimate  [95% confidence interval]** |
| --- | --- |
| Intercept | 3.747 [3.261 / 4.196] |
| **Habitat (Evergreen)** | **-0.860 [-1.715 / -0.006 ]** |
| **Habitat (Mixed)** | **-0.816 [-1.496 / -0.134]** |
| Day length | 0.026 [-0.022 / 0.075] |
| Precipitation | 0.318 [-0.022 / 0.656] |
| Species (Great tit) | 0.282 [-0.160 / 0.723] |

**Table S5:** Effect of latitude, longitude, habitat type (deciduous, mixed, evergreen), and species (great or blue tit) on the midpoint (Julian day), duration (days), and delay (days) of temperature windows during which temperature most strongly affects laying date. Models including interactions between species and both latitude and longitude. Significant terms (where 95% confidence intervals do not overlap with 0) are shown in bold. The blue tit is used as the reference category for the species term. Deciduous is used as the reference category for the habitat type term.

| **Predictor variable** | **Parameter estimate  [95% confidence interval]** |
| --- | --- |
| **Midpoint** |  |
| Intercept | 5.099 [-47.735 / 52.534] |
| **Latitude** | **1.499 [0.564 / 2.543]** |
| Longitude | 0.406 [-0.201 / 0.970] |
| Species (Great tit) | -10.972 [-52.392 / 30.518] |
| Habitat (Evergreen) | 9.578 [-2.443 / 22.078] |
| Habitat (Mixed) | -0.296 [-9.635 / 8.702] |
| Species (Great tit):Latitude | 0.384 [-0.464 / 1.189] |
| **Species (Great tit):Longitude** | **-0.523 [-0.965 / -0.048]** |
| **Duration** |  |
| Intercept | 38.320 [-45.420 / 115.942] |
| Latitude | 0.401 [-1.148 / 2.083] |
| Longitude | -0.675 [-1.694 / 0.304] |
| Species (Great tit) | 33.673 [-63.604 / 135.169] |
| Habitat (Evergreen) | -7.985 [-24.364 / 9.050] |
| Habitat (Mixed) | -9.203 [-22.245 / 2.672] |
| Species (Great tit):Latitude | -0.736 [-2.737 / 1.194] |
| Species (Great tit):Longitude | 0.707 [-0.359 / 1.831] |
| **Delay** |  |
| **Intercept** | 41.946 [7.26 / 74.692] |
| Latitude | -0.247 [-0.9 / 0.458] |
| Longitude | -0.506 [-0.925 / -0.099] |
| Species (Great tit) | -6.817 [-42.627 / 31.179] |
| Habitat (Evergreen) | 2.052 [-5.143 / 9.814] |
| Habitat (Mixed) | 1.584 [-4.363 / 7.095] |
| Species (Great tit):Latitude | 0.026 [-0.737 / 0.742] |
| Species (Great tit):Longitude | 0.352 [-0.047 / 0.779] |

**Table S6:** Effect of species (great or blue tit), precipitation, habitat type (deciduous, mixed, evergreen) and day length (minutes) on the strength of phenological sensitivity (days/^o^C). Models including interactions between species and all other fixed effects terms. Significant terms (where 95% confidence intervals do not overlap with 0) are shown in bold. The blue tit is used as the reference category for the species term and deciduous habitat is the reference category for habitat type.

| **Predictor variable** | **Parameter estimate  [95% confidence interval]** |
| --- | --- |
| **Intercept** | **3.814 [3.26 / 4.304]** |
| Species (Great tit) | 0.194 [-0.449 / 0.856] |
| Habitat (Evergreen) | -1.106 [-2.347 / 0.148] |
| Habitat (Mixed) | -0.855 [-1.888 / 0.108] |
| Day Length | 0.028 [-0.038 / 0.091] |
| **Precipitation** | **0.456 [0.073 / 0.841]** |
| Species (Great tit):Habitat (Evergreen) | 0.369 [-1.243 / 1.902] |
| Species (Great tit):Habitat (Mixed) | 0.046 [-1.248 / 1.281] |
| Species (Great tit):Day Length | -0.003 [-0.081 / 0.077] |
| Species (Great tit):Precipitation | -0.093 [-0.753 / 0.573] |
