## Supplementary material 2 for "Bird populations most exposed to climate change are less responsive to climatic variation"

**Supplement Material 2: Explanation on how a covariance between the exposure and sensitivity affects the mean and variation in expected phenological advancement**

We found a negative covariance in the exposure and sensitivity of great and blue tit populations: the populations with the strongest exposure tended to be the least sensitive. Intuitively one might expect that a negative covariance will reduce intra-specific variation in expected phenological advancement compared to a species in which there is no covariance between the sensitivity and exposure, while a positive covariance will increase the intra-specific variation. Here we explore this intuition formally, by mathematically describing how any among-population covariance between sensitivity and exposure will affect the mean and variance in expected phenological advancement for the entire species.

We will compare two species that have the same mean and variance in sensitivity (X) and exposure (Y) but where one species has a negative covariance between sensitivity and exposure (COV(X,Y)<0), while the other species shows no covariance between sensitivity and exposure (COV(X,Y)=0). We are interested in how COV(X,Y) affects the mean (expectation) and variance in expected phenological advancement (XY, the product of X and Y) for the species as a whole.

*How covariance between sensitivity and exposure affects the mean expected phenological advancement*

Probability theory tells us that the expectation (E) of the product of two independent variables X and Y is given by:

$${E[X*Y]}_{i}=E(X)*E(Y)$$

Probability theory also tells us that the expectation of the product of two dependent variables is given by:

$${E[X*Y]}_{d}=E(X)*E(Y)+COV(X,Y)$$

From this it can be seen that a negative covariance lowers the expected value of a species mean phenological advancement if we compare it to a species in which sensitivity X and exposure Y are independent, since:

$${E[X*Y]}_{d}-{E[X*Y]}_{i}=COV(X,Y)$$

Furthermore, the relative contribution of any covariance between sensitivity and exposure to the observed expected phenological advancement ($R_{c\to{E\left[ X*Y \right]}_{d}}$) can be quantified as:

$$R_{c\to{E\left[ X*Y \right]}_{d}}=\frac{{E\left[ X*Y \right]}_{d} - {E\left[ X*Y \right]}_{i}}{{E\left[ X*Y \right]}_{d}}=\frac{COV(X,Y)}{E(X)*E(Y)+ COV(X,Y)}$$

In our dataset (Table S1) we found that mean exposure is $E\left( X \right)=0.0354$˚C/year, mean sensitivity is $E\left( Y \right)=3.614$days advancement/˚C and the covariance $\mathrm{COV}\left( X,Y \right)=0.00361$. This means that $R_{c\to{E\left[ X*Y \right]}_{d}}=-0.029$ which tells us that the covariance between sensitivity and exposure lowered the mean expected phenological advancement by only 2.9% compared to a situation with no covariance.

Although we have shown above that the mean expected phenological advancement across all populations was little affected by the covariance, we emphasize that for calculating the expected advancement of a single population with a given sensitivity it can make a huge difference whether one uses the population specific exposure or the exposure averaged across all populations.

*How covariance between sensitivity and exposure affects the intraspecific variation in expected phenological advancement*

Probability theory tells us that the variance (V) of the product of two independent variables is given by:

$${V[X*Y]}_{i}=V(X)*V(Y)+V(X)*{E(Y)}^{2}+V(Y)*{E(X)}^{2}$$

Probability theory also tells us that the variance (V) of the product of two dependent variables is given by:

$${V\left[ X*Y \right]}_{d}=V\left( X \right)*V\left( Y \right)+V\left( X \right)*{E\left( Y \right)}^{2}+V\left( Y \right)*{E\left( X \right)}^{2}$$

$$+{E\left( X \right)}^{2}*{E\left( Y \right)}^{2}+COV(X^{2},Y^{2})-{(COV(X,Y)+E(X)*E(Y))}^{2}$$

The influence of a negative covariance on the intraspecific variance of a species expected phenological advancement can thus be quantified as:

$${V\left[ X*Y \right]}_{d}-{V[X*Y]}_{i}={E\left( X \right)}^{2}*{E\left( Y \right)}^{2}+COV(X^{2},Y^{2})-{(COV(X,Y)+E(X)*E(Y))}^{2}$$

From this it can be seen that the contribution of any covariance to the intraspecific variance is more complex, as it also depends on other factors (i.e. on $COV(X^{2},Y^{2})$, $E\left( X \right)$ and $E(Y)$). Consequently, a negative covariance does not in all cases reduce the intra-specific variability in phenological advancement.

Finally, the relative contribution of any covariance between sensitivity and exposure to the observed variance in phenological advancement (R_c_) can be quantified as:

$$R_{c\to{V\left[ X*Y \right]}_{d}}=\frac{{V\left[ X*Y \right]}_{d} - {V\left[ X*Y \right]}_{i}}{{V\left[ X*Y \right]}_{d}}=$$

$$\frac{{E\left( X \right)}^{2}*{E\left( Y \right)}^{2}+COV(X^{2},Y^{2})- {( COV(X,Y)+E(X)*E(Y) )}^{2}}{V(X)*V(Y)+V(X)*{E(Y)}^{2}+ V(Y)*{E(X)}^{2} +{E\left( X \right)}^{2}*{E\left( Y \right)}^{2}+ COV(X^{2},Y^{2})- {( COV(X,Y)+E(X)*E(Y) )}^{2}}$$

In our dataset (Table S1) we found that variance in exposure is $V\left( X \right)=0.00749$(˚C/year)^2^, variance in sensitivity is $V\left( Y \right)=1.1867$(days advancement / ˚C)^2^ and the covariance $\mathrm{COV}\left( X^{2},Y^{2} \right)=-0.00332$ . As a result$R_{c\to{V\left[ X*Y \right]}_{d}}=-0.2367$ and thus the covariance lowered the intraspecific variance in phenological advancement by 23.7% compared to a situation in which there was no covariance.
